# Immune Regulation of Intestinal Stem Cell Proliferation and Differentiation in *Drosophila*

**DOI:** 10.1101/721662

**Authors:** Minjeong Shin, Meghan Ferguson, Reegan J. Willms, Lena O. Jones, Kristina Petkau, Edan Foley

## Abstract

Intestinal progenitor cells integrate signals from their niche, and from the gut lumen, to divide and differentiate at a rate that maintains an epithelial barrier to microbial invasion of the host interior. Despite the importance of evolutionarily conserved innate immune defenses to maintain stable host-microbiota relationships, we know little about specific contributions of stem cell immunity to gut homeostasis. We used the *Drosophila* model to determine the consequences of compromised intestinal stem cell immune activity for epithelial homeostasis. We showed that loss of stem cell immunity greatly impacted growth and renewal in the adult gut. In particular, we noticed that inhibition of stem cell immunity impeded key growth and differentiation events in the progenitor cell compartment leading to a gradual loss of stem cell numbers with age, and an impaired differentiation of mature enteroendocrine cells. Our results highlight the importance of immune signaling in the stem cell population for epithelial function in the adult gut.

**HIGHLIGHTS:**

- The TNFR-like Immune Deficiency (IMD) pathway is active in *Drosophila* intestinal progenitor cells.
- Inhibition of IMD in progenitors impairs progenitor cell proliferation.
- Blocking progenitor cell IMD negatively affects generation of mature epithelial cells.

## INTRODUCTION

The intestinal epithelium is an important contact point between an animal and their environment. Collectively, intestinal epithelial cells regulate nutrient acquisition, microbiota tolerance, immune education, and pathogen elimination, and disruptions to epithelial homeostasis are linked to inflammatory diseases and cancers. As the epithelium consists of a heterogenous population of specialized cell types, it is essential that we understand the mechanisms by which individual lineages regulate intestinal cell proliferation, differentiation, and renewal.

Intestinal epithelial cell (IEC) lineages vary by animal, but data from *Drosophila*, zebrafish, mice, and humans indicate broad evolutionarily conservation of intestinal cell-type composition (Brugman, 2016; Buchon et al., 2013a; Lickwar et al., 2017; Miguel-Aliaga et al., 2018; Nguyen et al., 2015; Wallace et al., 2005). Typically, the epithelium is maintained by proliferative, multipotent Intestinal Stem Cells (ISCs) that self-renew, and generate all differentiated epithelial cell types. Most terminally differentiated cells are columnar enterocytes, a cell type specialized in capture and digestion of lumenal nutrients. Secretory cell-type complexity varies from animal to animal. In flies, the secretory lineage appears to consist solely of hormone-producing enteroendocrine cells. In contrast, fish and mammals have mucus-secreting goblet cells in addition to the enteroendocrine population, and mammals also have long-lived, antimicrobial peptide-producing Paneth cells that neighbor ISCs in basal crypts. ISCs integrate cues from their niche to divide and differentiate at a rate that replenishes dying epithelial cells, effectively maintaining gut function, and barrier integrity. Notch, Epidermal Growth Factor (EGF), Wnt and Bone Morphogenetic Protein (BMP) signal transduction pathways are particularly important regulators of ISC proliferation and differentiation in vertebrates and invertebrates (Spit et al., 2018). More recent studies uncovered roles for immune signaling in ISC survival, growth and differentiation. For example, vertebrate ISCs express MHC class II molecules, and presentation of self-peptides by ISCs appears to be a critical aspect of intestinal invasion and epithelial destruction in graft-versus host disease (Biton et al., 2018; Fu et al., 2019; Takashima et al., 2019). Likewise, ISCs are enriched for expression of the germline-encoded peptidoglycan receptor NOD2 (Nigro et al., 2014). NOD2 protects ISCs against reactive oxygen species toxicity (Levy et al., 2020), and mutations in NOD2 are associated with Crohn’s disease and intestinal tumorigenesis (Couturier-Maillard et al., 2013). Despite established requirements for immune signaling pathways in the maintenance of intestinal health, it is unclear if innate defenses act specifically in progenitors to regulate epithelial homeostasis. We consider this an important knowledge gap given the central role of intestinal progenitors in building and maintaining the entire epithelium.

*Drosophila melanogaster* are widely used to characterize intestinal epithelial immunity and homeostasis. The adult fly intestine is a pseudostratified epithelium that is maintained by multipotent ISCs (Micchelli and Perrimon, 2006; Ohlstein and Spradling, 2006). The majority of *Drosophila* ISC divisions are asymmetric, producing a new ISC, and a transient cell type that generates terminally differentiated epithelial cells. In most cases, ISC divisions generate a post-mitotic enteroblast that differentiates as an enterocyte in response to Notch signals (Bardin et al., 2010; Guo and Ohlstein, 2015; Ohlstein and Spradling, 2007). Collectively, ISCs and enteroblasts are classified as the intestinal progenitor compartment in flies. Alternatively, in the absence of cues from Notch, ISCs transition through a pre-enteroendocrine state to generate mature enteroendocrine cells that can be sub-classified into functional groups based on intestinal localization, and hormone expression patterns (Biteau and Jasper, 2014; Guo and Ohlstein, 2015; Zeng and Hou, 2015). In the fly gut, bacterial diaminopimelic acid-type peptidoglycan (PGN) activates immune responses via the Immune Deficiency (IMD) pathway, a germline-encoded defense with similarities to vertebrate Tumor Necrosis Factor Receptor signaling (Buchon et al., 2009b, 2013a; Myllymäki et al., 2014). Detection of extracellular polymeric PGN by the PGRP-LC receptor, or intracellular monomeric PGN by the related PGRP-LE receptor converges on a proximal signaling complex that includes the Imd protein, the adaptor protein FADD, and the Caspase-8 homolog, Dredd. Dredd proteolytically removes thirty N-terminal amino acids from Imd, initiating molecular events that activate c-Jun N-terminal Kinase, and the p100/105 NF-kB ortholog Relish (Rel) (Leulier et al., 2000; Stoven et al., 2003; Stöven et al., 2000). Thus, Dredd-mediated processing of Imd is essential for IMD pathway activation, and expression of a non-cleavable Imd variant (ImdD30A) blocks host responses to PGN in cell culture and *in vivo* (Kim et al., 2014; Paquette et al., 2010). In the fly intestine, Rel and c-Jun N-terminal Kinase initiate transcriptional responses that include regionalized expression of antimicrobial peptides, regulators of metabolism, and genes associated with growth and differentiation (Broderick et al., 2014; Buchon et al., 2009b, 2009a; Dutta et al., 2015; Hung et al., 2020). Earlier work uncovered significant differences between the responses of mature epithelial cell-types to IMD activation. For example, infection-dependent activation of IMD in enterocytes results in antimicrobial peptide expression and extrusion of damaged cells (Buchon et al., 2009b; Dutta et al., 2015; Zhai et al., 2018). In contrast, activation of IMD in enteroendocrine cells by commensal-derived lactate modifies lipid metabolism in neighboring enterocytes (Kamareddine et al., 2018). Notably, genomic studies demonstrated expression of IMD pathway components in intestinal progenitor cells (Dutta et al., 2015; Hung et al., 2020). However, the contributions of progenitor-specific IMD activity to intestinal homeostasis are unexplored.

We took advantage of the genetic accessibility of flies to ask if progenitor-specific IMD activity affects intestinal homeostasis in adult *Drosophila*. Specifically, we used genomic and physiological assays to determine the consequences of blocking IMD in intestinal progenitors. We found that inhibition of progenitor cell IMD had significant effects on ISC proliferation, progenitor compartment composition, and generation of mature enteroendocrine cells. As germline-encoded immune responses are known modifiers of vertebrate intestinal epithelial growth, we believe our findings are of general relevance to understanding how host immune responses control stem cell function in the intestine.

## RESULTS

### Inhibition of IMD Affects the Intestinal Progenitor Cell Transcriptome

In a single-cell RNA sequencing profile of adult female *Drosophila* intestines, we identified 620 cells that expressed canonical progenitor markers such as *esg*, *Dl*, and *N* (Figure S1). Within the progenitor population, we observed enriched expression of key IMD pathway components, including the peptidoglycan sensor *pgrp-lc*, the NF-kB transcription factor *relish*, and the IMD pathway target *pirk* (Figure 1A, B). Our data match earlier reports of IMD pathway gene expression in progenitors (Dutta et al., 2015; Hung et al., 2020), and raise the possibility that immune signals contribute to progenitor cell function in the fly gut.

**Figure 1.**
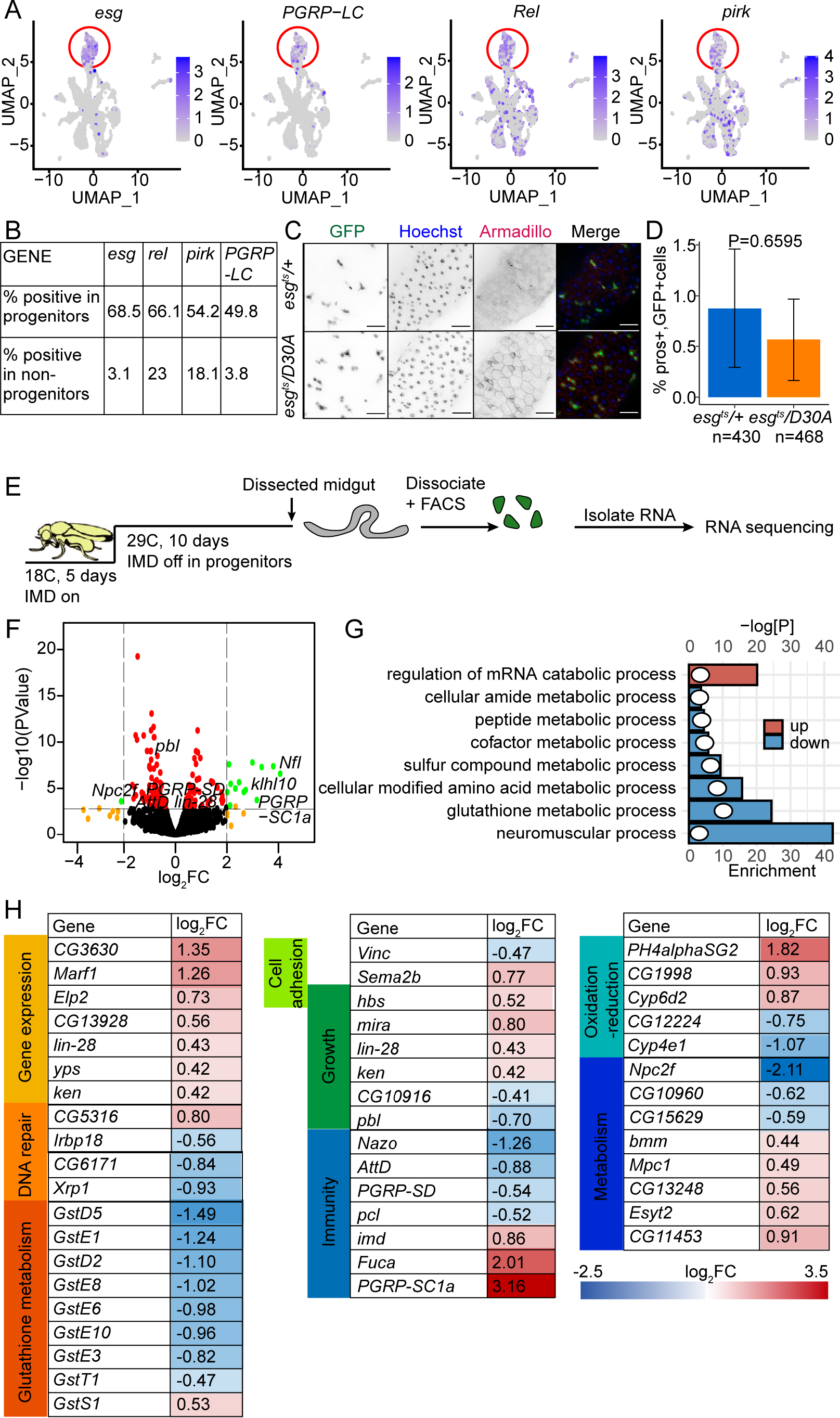
IMD Regulates the intestinal progenitor cell transcriptome. (A) Uniform Manifold Approximation and Projection (UMAP) plot of intestinal epithelial cells isolated from ten-day-old *esg^ts^* flies showing expression of the progenitor marker *esg*, and the IMD pathway components *PGRP-LC*, *Rel*, and *pirk*. Progenitors are circled in red. (B) Quantification of the percentage of progenitors and non-progenitors that express the indicated genes. (C) Visualization of GFP, DNA (Hoechst) and the beta-catenin ortholog in intestines of ten-day-old *esg^ts^*/*+* and *esg^ts^*/*UAS-imdD30A (esg^ts^*/*D30A*) flies. (D) Quantification of the percentage of GFP-positive cells that express the enteroendocrine cell marker *prospero* in *esg^ts^*/*+* (n=20) and *esg^ts^*/*D30A* (n=22) flies. (E) Schematic representation of an experimental strategy to quantify gene expression in progenitor cells purified from *esg^ts^*/*+* and *esg^ts^*/*D30A* flies. Each experiment was performed in triplicate. (F) Volcano plot showing relative changes (x-axis) and significance (y-axis) in gene expression of purified progenitors from *esg^ts^*/*D30A* flies compared to age-matched *esg^ts^*/*+* controls. Genes are color coded to indicate significance and relative gene expression changes. (G) Gene Ontology analysis of processes significantly affected by inhibition of IMD in progenitors. Column size indicates the degree of enrichment for each term, and dots indicate the log-transformed significance of the respective enrichment. (H) Representative sample of genes with affected expression upon inhibition of IMD in progenitors.

To test if IMD is functionally active in progenitors, we used the *esgGAL4*, *GAL80^ts^*, *UASGFP* (*esg^ts^*) fly line to express a dominant inhibitory IMD protein (ImdD30A) in *Drosophila* progenitors (*esg^ts^/D30A*) for ten days. Notably, blocking IMD in progenitors did not prevent infection-mediated expression of IMD-responsive antimicrobial peptides in enterocytes, demonstrating that inhibition of progenitor cell IMD does not affect IMD activity in differentiated progeny (Figure S2). We then asked if blocking IMD has direct effects on the progenitor population. Intestines of *esg^ts^/D30A* and control *esg^ts^/+* flies had similar distributions of small, GFP-positive cells (Figure 1C) that rarely expressed the enteroendocrine cell marker, *prospero* (Figure 1D), confirming that GFP exclusively marked progenitors in both lines. To measure effects of IMD inhibition on progenitors, we performed RNAseq analysis of gene expression in FACS-purified GFP-positive cells from *esg^ts^/D30A* and *esg^ts^/+* flies (Figure 1E). We found that inactivation of IMD disrupted expression of 154 genes in progenitors (Figure 1F, table S1), including key IMD pathway regulators (*pgrp-sd*, *pgrpsc1a*), glutathione metabolism genes required for detoxification of xenobiotic substances, and 24 genes known to respond to the commensal microbiome (Broderick et al., 2014) (Figure 1F-H). Notably, the impacts of IMD inhibition extended beyond conventional antimicrobial responses and included diminished expression of genes associated with stem cell growth, and adhesion to the niche, such as the growth regulator *Xrp1*, the asymmetric cell division regulator *miranda* (*mira*), and the effector of extracellular matrix adhesion *Vinculin* (*Vinc*, Figure 1F-H), suggesting potential growth-regulatory roles for IMD in progenitors.

### Blocking IMD in Progenitors Inhibits ISC Division

As IMD inhibition affected the progenitor cell transcriptome, we asked if IMD also affects progenitor cell homeostasis. Specifically, we measured ISC mitoses by quantifying phospho-histone H3; the percentage of midgut epithelial cells that expressed the progenitor marker *esg*; and the percentage of progenitors that expressed the ISC marker *Delta* (*Dl*) in five and thirty-day-old *esg^ts^/D30A* and *esg^ts^/+* intestines. In young *esg^ts^/+* intestines, we observed few ISC divisions (Figure 2A), approximately 20% *esg*+ cells in the posterior midgut (Figure 2B), and 40% Dl+ ISCs per progenitor (Figure 2C). Consistent with reports of age-related decline in gut function, we detected significantly increased numbers of ISC divisions, *esg*+ cells, and Dl+ cells in aged *esg^ts^/+* intestines relative to their five-day-old counterparts (Figure 2A-C). In contrast, we did not detect age-dependent increases in mitoses, progenitor number, or ISC numbers in thirty-day-old *esg^ts^/D30A* intestines compared to their five-day-old counterparts (Figure 2A-C). Instead, thirty-day-old *esg^ts^/D30A* intestines were characterized by significantly fewer mitoses, Dl+ ISCs, and progenitors than thirty-day-old *esg^ts^/+* control flies. Importantly, these results are not an artifact of *imdD30A* expression, as progenitor-specific, RNAi-mediated depletion of the IMD pathway adaptor FADD also caused a significant decline in the amounts of Dl-positive stem cells among intestinal progenitors of thirty-day-old flies (Figure 2D). Furthermore, we found that progenitor-specific inactivation of *relish* significantly impaired generation of mitotic clones in the posterior midgut (Figure 2E), confirming that genetic inhibition of IMD blocks intestinal epithelial proliferation. ISC-specific inactivation of IMD was sufficient to block proliferation (Figure 2F), whereas EB-specific inhibition of IMD had no effect (Figure 2G), suggesting a cell-autonomous role for IMD in controlling the rate of ISC division. Finally, we discovered that inhibition of the PGN sensors *PGRP-LC* or *PGRP-LE* (Figure 2H-I) was sufficient to inhibit progenitor cell proliferation. Collectively, our data indicate that inactivation of IMD in progenitors significantly impairs age-dependent accumulation of mitotically active progenitors in the adult midgut. We consider these findings particularly interesting, as increased epithelial immune responses are a hallmark of the aging intestine (Broderick et al., 2014; Buchon et al., 2009a; Guo et al., 2014; Ren et al., 2007).

**Figure 2:**
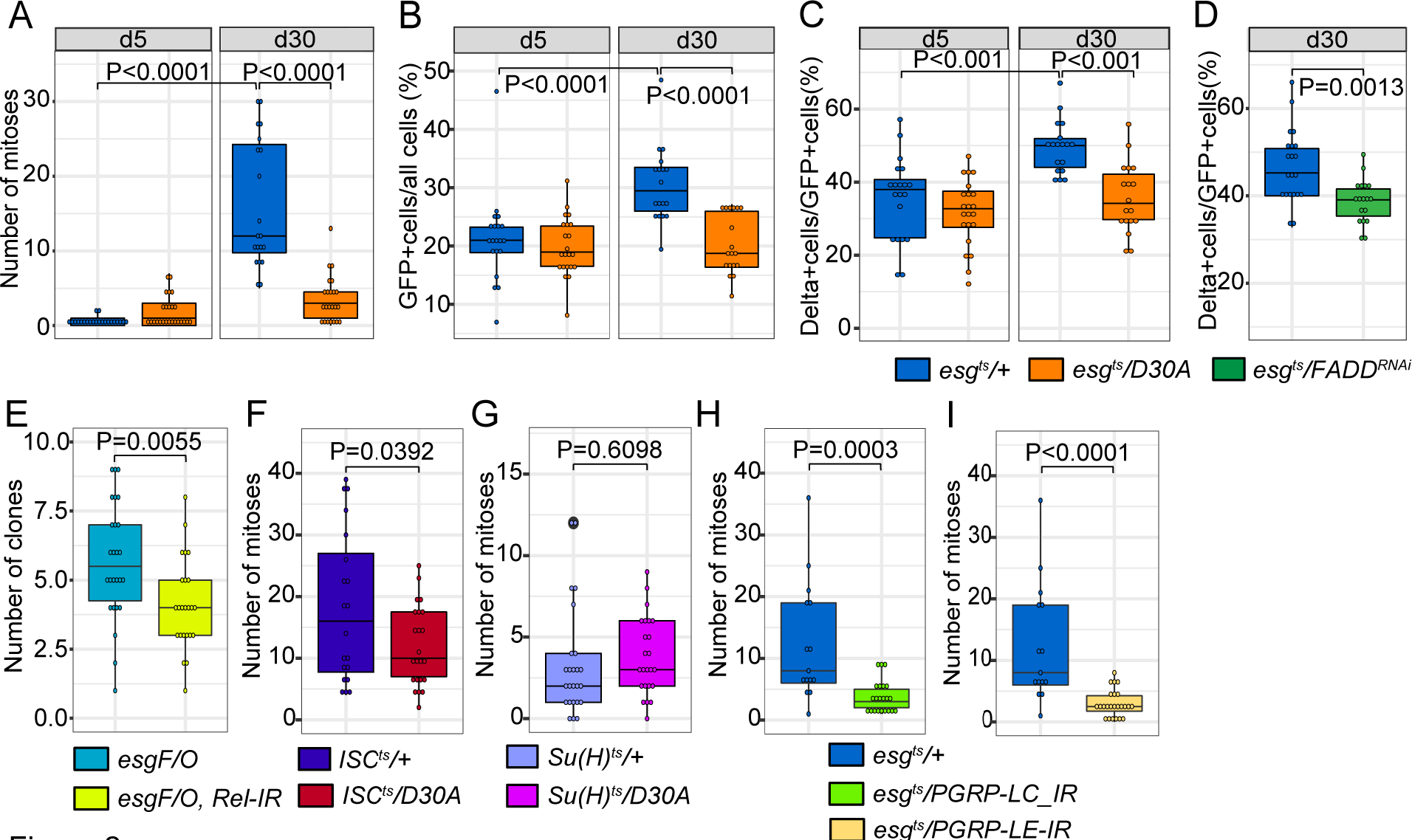
Progenitor cell IMD activity modifies stem cell proliferation. (A) Quantification of mitoses per gut in intestines from *esg^ts^*/*+* and *esg^ts^*/*D30A* flies of the indicated ages. (B) Percentage of intestinal epithelial cells that express the progenitor marker *esg* in *esg^ts^*/*+* and *esg^ts^*/*D30A* flies. (C) Percentage of progenitors that express the stem cell marker Delta in *esg^ts^*/*+* and *esg^ts^*/*D30A* flies. For A-C n=20 at d5, and 18 at d30 in *esg^ts^/+* flies and n=22 at d5, and 18 at d30 in *esg^ts^*/*D30A* flies. (D) Percentage of progenitors that express the stem cell marker Delta in *esg^ts^*/*+* (n=21) and *esg^ts^*/*FADD^RNAi^* (n=18) flies. (E) Quantification of GFP-marked mitotic clones in the posterior midgut of *esg^F/O^* (n=26) and *esg^F/O^*, *rel-IR* (n=24) flies nine days after marking of mitotic clones. (F) Quantification of mitoses per gut in *ISC^ts^/+* ( n=21) and *ISC^ts^/D30A* (n=24) 27 days flies. (G) Quantification of mitoses per gut in 27-day-old *Su(H)^ts^*/+ ( n=25) and *Su(H)^ts^/D30A* (n=24) flies. (H-I) Quantification of mitoses per gut in 27-day-old *esg^ts^*/*+* ( n=15), *esg^ts^*/*PGRP-LC-IR* (n=22) (H) and *esg^ts^*/*PGRP-LE-IR* (I) flies. Statistical significance for A-C was calculated using an ANOVA followed by pairwise Tukey comparisons, and significance for D-I was calculated using a Student’s t-test.

### Single Cell Transcriptional Profiles Uncover Impacts of Progenitor Cell IMD on the Intestinal Epithelium

Similar to vertebrates, the *Drosophila* intestine is a highly heterogenous tissue. Multipotent stem cells generate distinct epithelial lineages that control nutrient acquisition, hormone production, and responses to intestinal microbes in a regionally specialized fashion. Thus, although our data implicate IMD in progenitor cell division, we do not yet understand the consequences of blocking IMD in progenitors for the entire intestine. To determine the effects of progenitor-specific IMD inhibition on all epithelial cell types, we resolved the transcriptomes of ten-day- old control *esg^ts^*/+ and *esg^ts^/D30A* intestines at the single cell level (Figure S3A-B). After excluding dead cells and doublets, we prepared single-cell RNA sequencing profiles of 3675 cells from *esg^ts^*/+ intestines, and 3654 cells from *esg^ts^/D30A* intestines. Using unsupervised graph-based clustering of data from *esg^ts^*/+ intestines, we identified all cell types previously described in the adult gut, including progenitors that expressed growth and differentiation regulators; enteroendocrine cells that produced peptide hormones; and enterocytes dedicated to digestion (Figure S3). A more detailed examination of single cell transcriptomes from *esg^ts^*/+ intestines uncovered clear signs of specialization among the individual cell types. Specifically, we discovered regionalized and cell-type-specific expression patterns for regulators of metabolism (Figure S4), growth (Figure S5), differentiation (Figure S6), peptidoglycan sensing and scavenging (Figure S7), and oxidative stress responses (Figure S8). Thus, our profile of *esg^ts^*/+ guts accurately recapitulated known features of spatial and functional specialization within the fly gut (Dutta et al., 2015; Hung et al., 2020), providing a reliable control for analysis of intestines with impaired progenitor cell IMD.

To determine if blocking IMD in progenitors affects the identity or function of mature epithelial cell types, we used the integrated data analysis workflow in Seurat to identify cell type-specific differences in gene expression patterns between *esg^ts^*/+ and *esg^ts^/D30A* intestines. Unsupervised clustering of the integrated data resolved distinct clusters of progenitors, enterocytes, and enteroendocrine cells, as well as cardia, copper cells, an enterocyte-like cluster, a cluster of immature enterocytes, and three lineages of unknown function (Figure 3B). Comparisons between the two genotypes suggested mild effects of blocking IMD in progenitors on the generation of mature intestinal epithelial cells. For example, we noted fewer EC-like cells, and considerably more cardia in intestines from *esg^ts^/D30A* than in *esg^ts^/+* flies (Figure 3B). Furthermore, blocking IMD in progenitors had significant effects on expression of genes associated with critical regulatory functions in the gut. For example, intestines from *esg^ts^/D30A* flies were characterized by shifts in RNA processing and translation in enterocytes, diminished precursor metabolite generation in enteroendocrine cells, and increased expression of genes involved in autophagy, cell polarity, and adhesion in progenitors (Figure 3C). Notably, blocking IMD in progenitors did not affect expression of antimicrobial peptides, or peptidoglycan recognition proteins in differentiated enterocytes (Table S2), further arguing that expression of *imdD30A* in progenitor cells does not inhibit immune activity in progeny. Instead, we observed substantial effects of inhibiting IMD on expression of genes with essential roles in progenitor cell division and polarity (Figure 3D), including the Notch signaling modifier *Npc2f*, the Notch pathway target *E*(*spl*)*m3-HLH*, and *Snakeskin* (*Ssk*) a key regulator of intestinal stem cell activity (Figure 3E). Thus, and consistent with data presented in Figures 1 and 2, our results indicate that inhibition of IMD in progenitors has significant effects on progenitor cell homeostasis.

**Figure 3.**
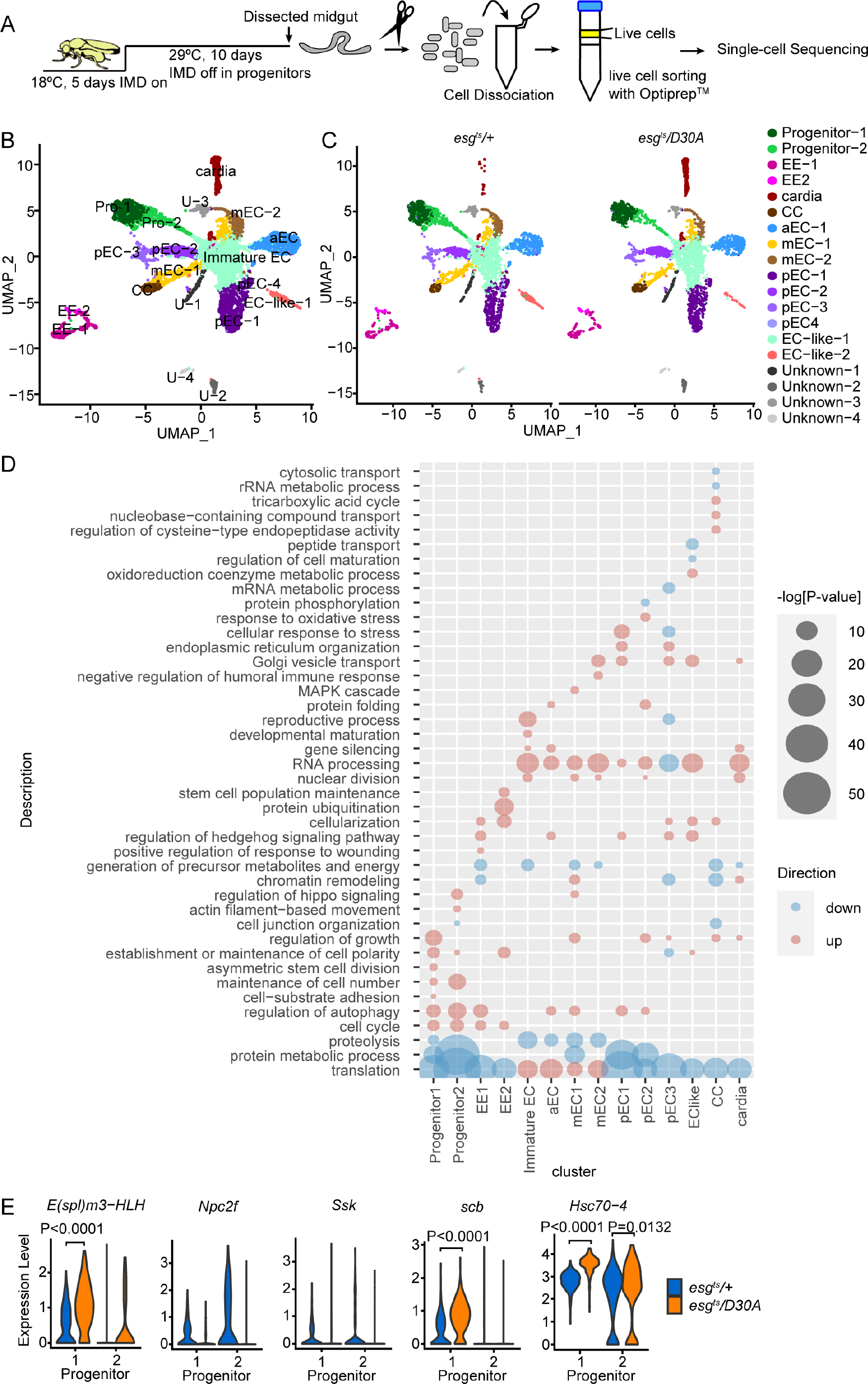
Inactivating IMD in progenitors affects transcriptional activity in all intestinal epithelial cell types. (A) Schematic representation of an experimental strategy for single-cell transcriptomic analysis of purified intestinal epithelial cells from *esg^ts^*/*+* and *esg^ts^*/*D30A* flies. (B) UMAP plot visualizing cell-types in integrated data from *esg^ts^*/*+* and *esg^ts^*/*D30A* flies based on the expression of marker genes. Cells are color coded by cell type. (C) The same data from panel B, split into the labeled genotypes. EE = enteroendocrine cells. CC = copper cells. EC = enterocytes subdivided according to anterior-posterior distribution along the intestine (a = anterior, m = middle, p = posterior). (D) Gene Ontology term analysis of cell-type specific processes significantly affected by progenitor-restricted inhibition of IMD. Bubble size indicates the log-transformed significance of the respective enrichments. Pink bubbles indicate enhanced terms, blue bubbles indicate underrepresented terms. € Representative violin plots of expression levels for the indicated genes in progenitors of *esg^ts^*/*+* and *esg^ts^*/*D30A* flies. P-values indicate significantly different expression levels. For *Npc2f* and *Snakeskin* (*Ssk*), no expression was observed in progenitors of *esg^ts^*/*D30A* flies.

As we believe our gene expression data are likely useful of value to the community outside the scope of the current study, we have deposited both sets on the Broad Institute Single Cell Portal (see Materials and Methods for further details).

### Inhibition of IMD Affects Developmental Trajectories Within the Progenitor Compartment

We were intrigued by our observation that blocking IMD in progenitors significantly affected expression of genes required for progenitor-niche interactions, and for progenitor differentiation. Therefore, we used Monocle to prepare developmental trajectories for *esg^ts^/+* and *esg^ts^/D30A* intestines in pseudotime. Analysis of the respective datasets successfully re-created developmental transitions from multipotent progenitors to differentiated lineages in both lines (Figure 4A, D). To directly examine effects of IMD on progenitors, we created subsets of the progenitor population for each genotype and analyzed gene expression in pseduotime for the respective subsets. Examination of the progenitor population from both genotypes revealed gene expression patterns characteristic of developmental transitions along a pseudotime trajectory (Figure 4B, E). For example, both progenitor populations were characterized by expression of the ISC marker *Dl* in early stages of pseudotime (Figure 4C, F). However, we found that IMD inhibition resulted in premature and prolonged pseudotime expression of the Notch targets *E(spl)m3-HLH* (Figure 4G-H) and *E(spl)malpha-BFM* (Figure 4I-J), the EGF receptor inhibitor *sprouty* (*sty*, Figure 4K-L), the EC fate regulator *klumpfuss* (*klu*, Figure 4M-N) and numerous markers of enterocyte maturation (Figure S9), indicating effects of progenitor cell IMD activity on enterocyte differentiation. To test if blocking IMD in progenitors has functional impacts on the transition from ISC to enteroblast, we monitored expression of fluorescent markers in *esgGAL4*, *UAS-CFP*, *Su(H)-GFP ; GAL80^ts^* flies that expressed ImdD30A. In these flies, ISCs are visible as CFP-positive cells (pseudo-colored as yellow), and enteroblasts are visible as CFP and GFP double- positive cells (pseudo-colored as magenta). Consistent with putative interactions between IMD and Notch signaling, we found that blocking IMD significantly increased the percentage of progenitors that expressed the enteroblast marker *Su*(*H*)*-GFP* (Figure 4O-P). Thus, in agreement with the loss of ISCs noted in *esg^ts^/D30A* intestines (Figure 2), our data argue that IMD activity influences progenitor cell composition in the fly intestinal epithelium.

**Figure 4.**
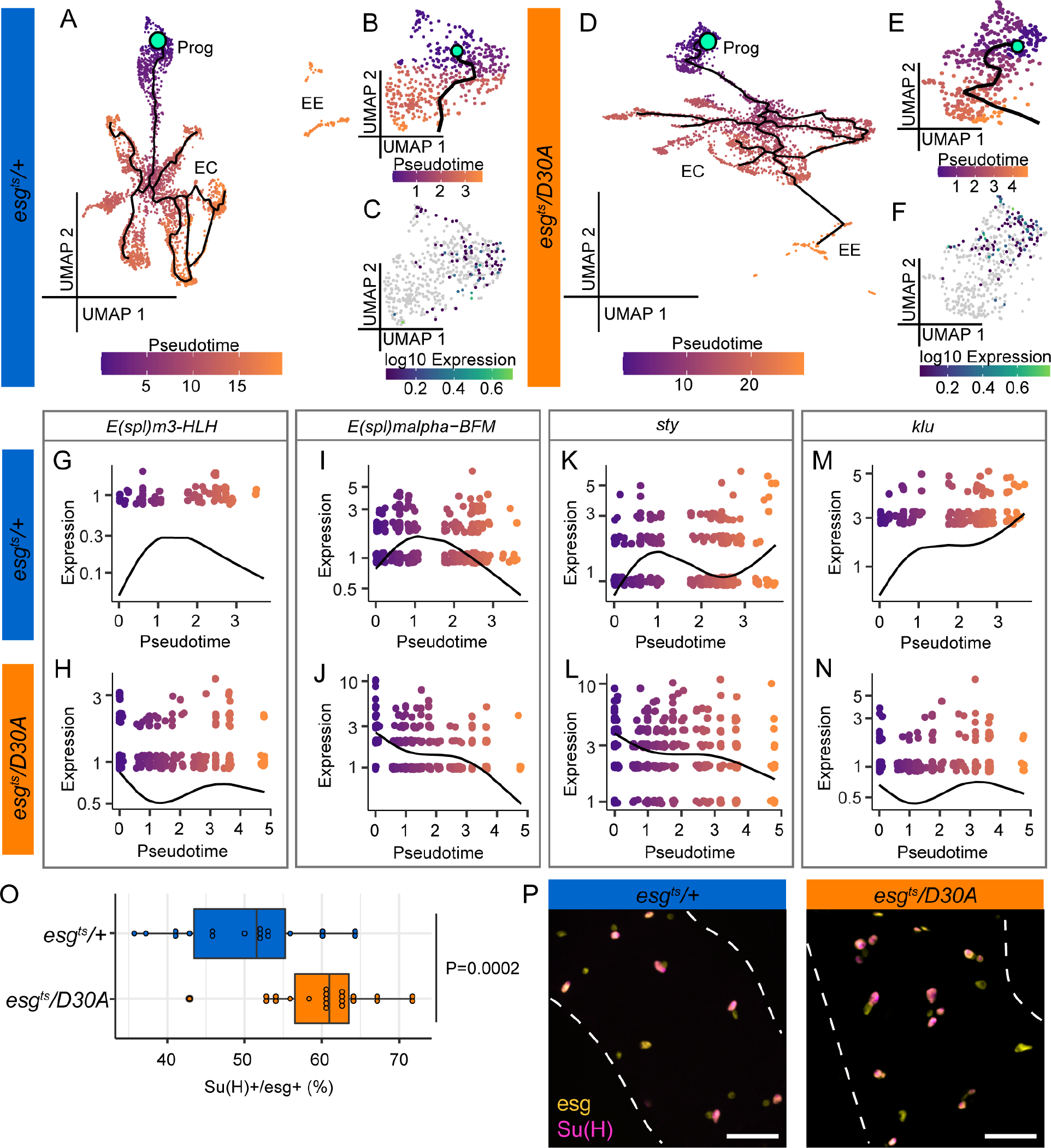
Inhibition of IMD affects developmental trajectories within the progenitor compartment. Single cell datasets from Seurat for each individual genotype were loaded into Monocle3 and pseudotime analysis was performed on midgut epithelial cells. (A) *esg^ts^/+* midguts with wild-type progenitors and (D) *esg^ts^/D30A* midguts with IMD-deficient progenitors. Mint green circles denote the root node and beginning of the intestinal trajectories. Dark purple marks cells at the beginning of pseudotime while orange marks cells late in pseudotime. Black lines show trajectories. Prog = Progenitors, EC = Enterocytes, EE = enteroendocrine cells. (B) and (E) show pseudotime within progenitor subsets of (A) and (D) respectively. (C, F) *Delta (Dl*) expression patterns within *esg^ts^/+* progenitors (C) and *esg^ts^/D30A* progenitors (F). Grey dots are cells with no detectable expression. (G-N) Expression of Notch target genes *E(spl)m3-HLH, E(spl)malpha-BFM*, the EGF inhibitor *sprouty (sty*), and the EC fate regulator *klumpfuss* (*klu*) over pseudotime within progenitor subsets of the indicated genotypes. (O) Percent of *esg+* progenitors that are positive for the enteroblast marker *Su(H)+* in *esg^ts^, UAS-CFP, Su(H)-GFP/+* (n=18) and *esg^ts^, UAS-CFP, Su(H)-GFP /D30A* (n=22) posterior midguts 14 days after transgene expression. Significance found using Students t-test. (P) Representative images of intestines used to gather data for panel O.

### Inhibition of Progenitor Cell IMD Affects Generation of Mature Enteroendocrine Cells

IMD inhibition impaired ISC proliferation, diminished ISC numbers, and impacted cell-type composition within the *esg*+ progenitor compartment, suggesting possible effects of progenitor cell IMD on development of mature epithelial cells. To determine if inhibition of IMD in progenitors affects epithelial differentiation, we monitored the prospero-positive enteroendocrine (EE) cell population in *esg^ts^/+* and *esg^ts^/D30A* intestines. We focused on EE cells, as fly EE cells have been characterized to a single-cell resolution (Guo et al., 2019; Hung et al., 2020), permitting detailed comparisons between *esg^ts^/+* and *esg^ts^/D30A* intestines. *Drosophila* EE cells can be divided into subsets with distinct peptide hormone expression profiles that are stable during homeostasis or after recover from infection (Beehler-Evans and Micchelli, 2015). Therefore, we tested if inhibition of IMD in progenitors affected the representation of EE subsets in the intestine. Using unsupervised clustering of *prospero-*positive cells from *esg^ts^/+* and *esg^ts^/D30A* intestines, we found that EE cells clustered into five subsets in both genotypes (Figure 5A, B). In both genotypes, each EE subset had a signature hormone expression pattern (Figure 5C, D). In some cases, subset-restricted expression patterns were conserved between *esg^ts^/D30A* and *esg^ts^/+* EE cells. For example, matching an earlier characterization of EE subsets (Beehler-Evans and Micchelli, 2015), we found that cells from *esg^ts^/+* subset zero, and from *esg^ts^/D30A* subset two expressed *Tk* and *Dh31*. Likewise, *esg^ts^/+* subset three cells and *esg^ts^/D30A* subset one cells were characterized by enhanced expression of *NPF*, and partial expression of *Gbp5* and *CCAP*. In contrast, we did not detect a counterpart of *esg^ts^/D30A* subset zero cells in *esg^ts^/+* controls, and we saw minimal conservation of *esg^ts^/+* subset zero gene expression patterns in *esg^ts^/D30A* EE cells, suggesting functional differences between EE cells in *esg^ts^/D30A* flies compared to *esg^ts^/+* flies. When we classified EE cells based on the number of peptides they expressed, we noted further differences between *esg^ts^/+* and *esg^ts^/D30A* intestines. In particular, fewer EE cells expressed zero, or one peptide, in *esg^ts^/D30A* guts than in *esg^ts^/+* guts, and a greater proportion expressed two or more peptides (Figure 5E). Likewise, for thirteen of fourteen peptides examined, a greater percentage of *esg^ts^/D30A* EE cells expressed the respective peptide than *esg^ts^/+* controls (Figure 5F), indicating enhanced peptide expression in *esg^ts^/D30A* flies. To directly test the effects of blocking IMD in progenitors on peptide expression levels, we performed an RNAseq analysis of dissected midguts from *esg^ts^/D30A* and *esg^ts^/+* flies. With the exceptions of *ITP* and *Gpb5*, we found that blocking IMD in progenitors resulted in increased expression of the remaining twelve peptides (Figure 5G), confirming a link between IMD inhibition and peptide hormone expression. Finally, we quantified EE numbers in posterior midguts of *esg^ts^/+* and *esg^ts^/D30A* flies that we raised at 29°C for ten days. We found that inhibition of IMD in progenitors decreased the proportion of mature EE cells by roughly 20% relative to *esg^ts^/+* controls (Figure 5H). Combined, our results establish that inhibition of progenitor cell IMD disrupts peptide hormone expression patterns in mature EE cells, decreases the amount of total EE cells, and increases the expression of most peptide hormones, confirming a link between progenitor cell IMD activity and EE cell development.

**Figure 5.**
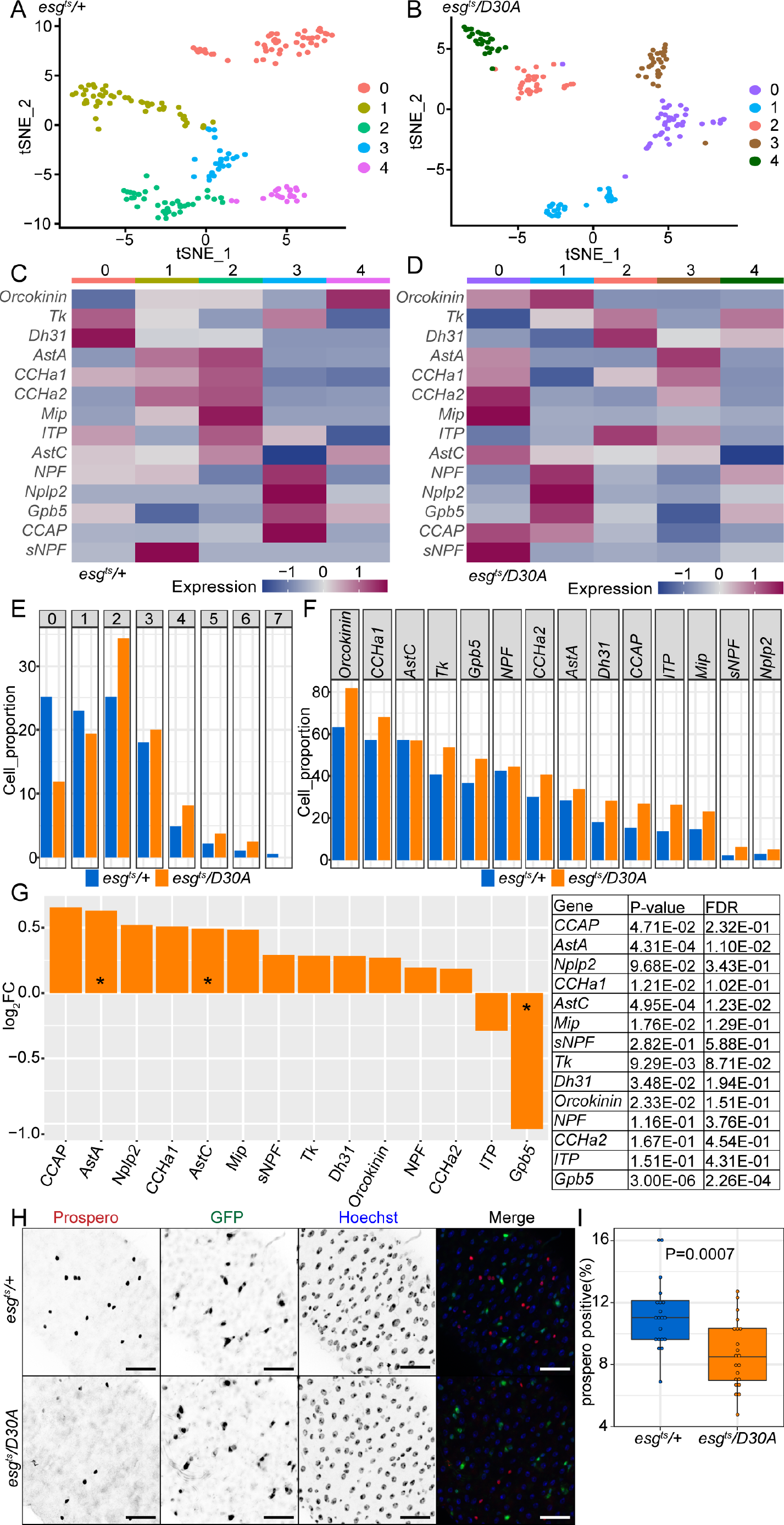
(A-B) tSNE plot visualizing subsets of *prospero*-positive enteroendocrine cells in *esg^ts^*/*+* (A) and *esg^ts^*/*D30A* intestines (B) based on the expression of marker genes. Cells are color coded by cell subset. (C-D) Heatmap showing relative expression of fourteen peptide hormones in each enteroendocrine cell subset in *esg^ts^*/*+* (C) and *esg^ts^*/*D30A* (D) intestines. (E) Quantification of the percentage of enteroendocrine cells that express the indicated numbers of peptide hormones. Genotypes are color coded as indicated. (F) Quantification of the percentage of enteroendocrine cells that express the indicated peptide hormone. Genotypes are color coded as indicated. (G) Quantification of the relative expression of each peptide hormone in isolated posterior midguts from *esg^ts^*/*D30A* flies relative to *esg^ts^*/*+* flies based on bulk RNAseq analysis. (H) Visualization of Prospero, GFP and DNA (Hoechst) in intestines of ten-day-old *esg^ts^*/*+* and *esg^ts^*/*D30A* flies. (I) Quantification of the percentage of intestinal epithelial cells that express the enteroendocrine cell marker prospero in the intestines of ten-day-old flies as indicated. Statistical significance was calculated using Students t-test. For *esg^ts^*/*+* flies n=20, and for *esg^ts^*/*D30A* flies n=21.

## DISCUSSION

Notch, BMP and WNT pathways regulate intestinal progenitor cell growth and differentiation in vertebrates and invertebrates (Barker, 2014; Casali and Batlle, 2009; Miguel-Aliaga et al., 2018; Sancho et al., 2015; Vooijs et al., 2011; Xu et al., 2011). In contrast, it is less clear what effects progenitor-specific activation of germline-encoded immune response pathways has on epithelial homeostasis. Several studies indicated survival and growth-regulatory effects of host immunity on intestinal progenitors. For example, the PGN pattern recognition receptor NOD2 is enriched in ISCs of mice (Nigro et al., 2014), and NOD2 protects ISCs from irradiation-induced cytotoxicity (Levy et al., 2020), while mutations in NOD2 are linked to Crohn’s disease (Hugot et al., 2001; Ogura et al., 2001). Likewise, TLR4 is expressed to higher levels in small and large intestinal crypts (Price et al., 2018), where its activation promotes crypt apoptosis and inhibits proliferation (Naito et al., 2017; Neal et al., 2012). In contrast, epithelium-wide activation of TLR4 promotes epithelial repair by activating EGF and JAK/STAT pathways in mice challenged with dextran sulfate sodium (Fukata et al., 2006; Hsu et al., 2010). In combination, these studies support roles for immune pathways in proliferative responses to cell-extrinsic challenges. However, we do not know if progenitor-specific immune activity impacts homeostatic growth and differentiation. We consider this an important question, as animal intestines contain dense microbial communities that promote growth and influence cell fate choices in intestinal epithelia of animals as divergent as flies, fish, and mice (Ferguson and Foley, 2021).

We measured growth and differentiation in midguts of adult *Drosophila* that we engineered to lack IMD activity in progenitors. The IMD pathway is highly similar to mammalian Tumor Necrosis Factor Receptor signaling, and IMD exerts broad regulatory effects on intestinal transcription (Broderick et al., 2014). In flies, IMD has context-dependent effects on ISC proliferation. IMD pathway mutants have elevated rates of mitoses that are driven by the microbiome (Buchon et al., 2009b; Guo et al., 2014; Paredes et al., 2011), but IMD is not required for the proliferative burst observed in flies challenged with *Ecc15* (Zhai et al., 2018). In contrast, infection with *Vibrio cholerae* blocks ISC proliferation in an IMD-dependent manner (Fast et al., 2020; Wang et al., 2013), whereas *Herpetomonas muscarum* induces IMD-dependent proliferation (Wang et al., 2019). In the absence of infectious agents, persistent activation of IMD in progenitors increases ISC division frequency and skews differentiation towards elevated numbers of enteroendocrine cells within the epithelium (Petkau et al., 2017). Similarly, overexpression of PGRP-LC in enterocytes induces Rel-dependent proliferation (Zhai et al., 2018). There are conflicting data on consequences of IMD inactivation in progenitors, with one study suggesting decreased proliferation (Wang et al., 2019), and a separate study indicating increased proliferation (Wang et al., 2013). We found that progenitor-specific inactivation of IMD diminished ISC proliferation, impaired age-dependent accumulation of *esg*-positive and Dl-positive progenitors, elevated the number of *Su*(*H*)-positive enteroblasts, and resulted in differentiation defects that included fewer enteroendocrine cells, and shifts in gene expression patterns associated with enterocyte subtypes. Thus, our work suggests that progenitor-specific immune activity contributes to epithelial homeostasis in flies and raises several questions about effects of progenitor cell IMD on the adult gut.

Which progenitor cell requires IMD to regulate epithelial differentiation? In flies, the progenitor compartment consists of undifferentiated ISCs, and post-mitotic enteroblasts that are committed to the enterocyte cell fate. Adult progenitors express IMD pathway components, and genomic studies, including data presented here, showed that ISCs and enteroblasts have highly similar gene expression profiles (Dutta et al., 2015; Hung et al., 2020), suggesting that both cell types are likely equally competent at IMD activation. ISCs are basally situated within the pseudostratified midgut epithelium, and are not expected to make frequent, direct contacts with the intestinal lumen. Enteroblasts are the apical daughters of ISC divisions that occur at oblique angles to the basement membrane. Thus, it seems more plausible that enteroblasts directly contact the lumen where they can detect PGN. However, it is important to note that gut-derived PGN is not strictly contained to the intestinal lumen in flies, or vertebrates. PGN crosses the intestinal epithelial barrier, even in the absence of detectable breaches, and several mechanisms are in place to prevent accumulation of PGN in the fly hemolymph (Capo et al., 2017; Gendrin et al., 2009; Paredes et al., 2011; Troha et al., 2019; Zaidman-Rémy et al., 2006). Thus, we cannot exclude the possibility that passive, or active, transport mechanisms allow diffusion of PGN across the gut barrier to ISCs, which in turn activate IMD and modulate differentiation responses in the hosts. In this regard, we consider it interesting that several vertebrate pattern recognition receptors have cell-type specific apicobasal distribution patterns. For instance, TLR4 is enriched apically in villi, and baso-laterally in crypts of the human colon (Fusunyan et al., 2001). Furthermore, apical stimulation of TLR9 promotes JNK activation, whereas basolateral stimulation of TLR9 leads to NF-kB activation, and IL-8 production (Lee et al., 2006). In future assays, it will be interesting to determine if basolateral detection of PGN influences fate choices within the progenitor compartment.

Is progenitor-specific IMD necessary to generate growth-regulatory enterocytes? Inhibition of progenitor cell IMD did not block IMD-dependent immune responses in enterocytes, confirming that the phenotypes reported here are not a consequence of ImdD30A perdurance in differentiated epithelia. However, we also found that inhibition of IMD in progenitors had consequences for epithelial differentiation, including effects on gene expression patterns within mature enterocytes. As enterocytes produce paracrine regulators of progenitor proliferation and differentiation, we cannot exclude the possibility that blocking IMD in progenitors disrupts enteroblast differentiation in a manner that modifies the ability of enterocytes to transduce growth and differentiation cues to progenitor cells. This may be particularly important in the context of severe epithelial damage, where secreted factors from dying enterocytes accelerate ISC proliferation to maintain the epithelial barrier and regenerate a mature gut. In this scenario, IMD activity in progenitors is important to establish homeostatic intercellular communications between enterocytes and the progenitor compartment, and loss of progenitor cell IMD interrupts a developmental loop between progenitors and enterocytes. Consistent with requirements for IMD in the control of epithelial differentiation, we noticed that blocking IMD in progenitors significantly affected the generation of mature enteroendocrine cells. Specifically, inhibition of progenitor cell IMD led to a decline in enteroendocrine numbers, but a general increase in the expression of peptide hormones, possibly as a compensatory mechanism. Notably, flies raised in an axenic environment have fewer enteroblasts and more enteroendocrine cells (Broderick et al., 2014), indicating non-overlapping contributions of progenitor cell immunity and gut microbes to developmental trajectories of ISCs.

Does IMD have regional effects on intestinal progenitor functions? Intestines are functionally specialized along the rostro-caudal axis, with distinct partitions governing various aspects of food digestion and absorption. IMD also displays clear signs of regional specialization. The foregut is characterized by enriched expression of PGRP- LC, and Rel regulates expression of chitin-binding proteins that contribute to peritrophic matrix construction (Buchon et al., 2009b; Neyen et al., 2012). In the midgut, PGN detection appears to rely primarily on PGRP-LE, particularly in the posterior midgut (Bosco-Drayon et al., 2012; Neyen et al., 2012). Activation of IMD in the anterior midgut results in expression of antimicrobial peptides that protect the fly from ingested microbes (Buchon et al., 2013b). In contrast, IMD activation leads to delamination of damaged cells in the midgut of flies infected with pathogenic bacteria, most notably in the R4 region of the posterior midgut (Zhai et al., 2018). In the posterior midgut, the transcription factor caudal prevents IMD-dependent antimicrobial peptide expression (Ryu et al., 2008). Instead, IMD induces expression of molecules that dampen immune signaling, including the PGRP- LC inhibitor *pirk*, and amidases that scavenge PGN (Bosco-Drayon et al., 2012). As a result, posterior midgut IMD activity establishes a tolerogenic environment for commensal bacteria. Notably, loss of IMD pathway inhibitors, or expression of PGRP-LC in enterocytes increases proliferation in the posterior midgut (Paredes et al., 2011; Zhai et al., 2018), raising the possibility that suppression of posterior midgut IMD activity is required to prevent excess proliferation in the absence of infection. With a large collection of genetic reagents, and accessible genomic methods, the fly is an excellent system to systematically characterize regional effects of immune responses on progenitor cell function. Given the evolutionary conservation of immune responses, we believe the findings reported in this study to be of relevance for understanding fundamental principles of immune-regulated intestinal homeostasis.

## MATERIALS AND METHODS

### Fly Husbandry

Flies were raised on corn meal medium (Nutri-Fly Bloomington formulation, https://bdsc.indiana.edu/information/recipes/bloomfood.html; Genesse Scientific) at 18°C or 25°C. All experimental flies were adult virgin females kept under a 12h:12h light:dark cycle and maintained at 18°C during collection then shifted to 29°C to express downstream genes as indicated. We used *w^1118^* as a wildtype strain and backcrossed *UAS-imdD30A* transgenic lines into the *w^1118^* background for eight generations prior to use and used standard husbandry methods to ensure that *esg^ts^* (*esg-GAL4*, *tub-GAL80^ts^*, *UAS-GFP*) flies had the same first and third chromosomes as our *w^1118^* line. Fly lines used in this study were: *w ; esg-GAL4,tubGAL80^ts^,UAS-GFP* (referred to as *esg^ts^*), *UAS-FADD^RNAi^*(VDRC ID# 7926), *w^1118^* (VDRC ID# 60000), *w;esg-GAL4,UAS-CFP, Su(H)-GFP;tubGal80^ts^* (*esg^ts^,UAS-CFP,Su(H)-GFP*), *GS 5961* (Mathur et al., 2010), *dpt-GFP*, *esg-GAL4, tubGAL80ts, UAS-GFP;UAS-flp, Act>CD2>GAL4* (referred to as *esg^F/O^*), *Esg[ts], Su(H) Gal80* (referred to as *ISC^ts^*), *Su(H)GBE-Gal4^ts^* (referred to as *Su(H)^ts^*), *PGRP-LE RNAi* (VDRC ID# 108199), *PGRP-LC RNAi* (VDRC ID# 101636), *Rel-RNAi* (VDRC ID# 49413), *40D-UAS* (control for VDRC KK lines, VDRC ID# 60101) . To induce GFP-marked mitotic clones using the *esg^F/O^* system, flies of the indicated genotype were raised at 18°C for three days after eclosion, shifted to 29°C for 16h, then raised at 25°C for an additional nine days.

### *Ecc15* Oral Infection

For oral infection with *Ecc15*, we incubated an overnight culture of *Ecc15* in LB (Difco^TM^ Luria Broth Base, Miller, 241420) supplemented with NaCl (4.75g Fisher Scientific, BP358-212 per 500mL of LB Broth base) at 29°C with shaking. Flies were starved (10 flies per vial) for 2 h before infection. *Ecc15* was pelleted at 1250g for 10 minutes at 4°C and supernatant decanted. The harvested bacterial pellet was re-suspended in residual LB and an equivalent volume of 5% sucrose in PBS. Flies were transferred into vials that contained a filter paper (Whatman^TM^, Grade 3, 23mm, 1003-323) soaked with 150ml of the *Ecc15* culture on top of standard corn meal medium. Flies were infected for 16h at 29° C with 12h:12h light:dark cycle. To activate the GeneSwitch (GS) system we added 100*μ*l RU486 (Mifepristone, M8046, Sigma) dissolved in 80% EtOH (5mg/ml) to the surface of standard fool and dried overnight prior to addition of flies. For controls, we added 100*μ*l of 80% EtOH to the surface of standard fool and dried overnight prior to addition of flies. Flies were raised on treated food for 48h prior to infection.

### Immunofluorescence

The number of PH-3 or Delta positive cells were analyzed with two-way ANOVA or unpaired Student’s t-tests. We used previously described immunofluorescence protocols to visualize posterior midguts (56). In brief, we used anti-phospho-histone H3 (PH3, 1:1000, Millipore (Upstate), 06-570) immunofluorescence to quantify mitoses in the midguts, and anti-Delta (1:100; Developmental Studies Hybridoma Bank (DSHB) C594.9B) immunofluorescence to quantify stem cells in the R4/R5 region of the posterior midguts of virgin female flies that we raised at 29 °C. We also used anti-prospero (1:100, DSHB), anti-armadillo (1:100, DSHB) as primary antibodies and Hoechst 33258 (1:500; Molecular Probes) for DNA staining. Secondary antibodies used: goat anti-mouse Alexa Fluor 568 (1:500; Invitrogen), goat anti-rabbit 488 (1:500; Invitrogen). Tissue was mounted in Fluoromount (Sigma- Aldrich F4680) and posterior midguts were visualized with a spinning disk confocal microscope (Quorum WaveFX; Quorum Technologies Inc.). Images were collected as Z-slices and processed with Fiji software to generate a Z- stacked image.

### Isolation of progenitor cell and RNA extraction

Progenitor cells were isolated by fluorescence activated cell sorting (FACS) as previously described by Dutta et al. (2013). Flies were raised at 29°C for 10 days. 100 fly guts per sample were dissected (malphighian tubules, foreguts, hindguts and crops removed) and placed into ice-cold 1XPBS/DEPC-treated water. Guts were dissociated with 1mg/ml of elastase at 27°C with periodic pipetting for 1h. GFP-positive progenitor cells were collected based on GFP fluorescence and size with a BD FACSAriaIII sorter. Cells were pelleted at 1200g for 5 minutes at 4°C and then resuspended in 500μl Trizol. Samples were stored at -80°C until all three biological replicates were collected. RNA was isolated via a standard Trizol-chloroform extraction and the RNA was sent on dry ice to the Lunenfeld- Tanenbaum Research Institute (Toronto, Canada) for library construction and sequencing. The sample quality was evaluated using Agilent Bioanalyzer 2100. TaKaRa SMART-Seq v4 Ultra Low Input RNA Kit for Sequencing was used to prepare full length cDNA. The quality and quantity of the purified cDNA was measure with Bioanalyzer and Qubit 2.0. Libraries were sequenced on the Illumina HiSeq3000 platform.

### Preparation of Single Cell Suspension for single cell RNAseq

Single-cell suspension preparation method were followed (Hung et al., 2018) with a few modifications. Flies were raised for 10 days at 29°C. Five guts were dissected at one time and moved to 1% BSA in PBS/DEPC-treated water. Once twenty-seven guts were dissected, we transferred the guts to 200ml 1XPBS/DEPC-treated water on the back side of a glass dissection plate (PYREX, 7220-85) and chopped with scissors. After mechanically fragmenting the tissue, it was transferred to a 1.5 ml tube containing 100ml 1XPBS/DEPC-treated water then enzymatically digested with elastase (final concentration 1mg/ml) at 27°C for 40 min with gentle pipetting every 10 min. The single cell suspension was pelleted at 300g for 15 min at 4°C and cell pellet resuspended in 200ml 0.04%BSA in 1XPBS/DEPC-treated water. The cell suspension was filtered through a 70*μ*m filter (300g for 1 min at 4°C).

Live cells were collected using OptiPrepTM Density Gradient Medium (SIGMA, D1556-250ML) using the OptiPrepTM Application Sheet C13 protocol. Briefly, a 40% (w/v) iodixanol working solution was prepared with 2 volumes of OptiPrepTM and 1 volume of 0.04 %BSA in 1XPBS/DEPC-treated water. This working solution was used to prepare a 22% (w/v) iodixanol solution in the same buffer. One volume of working solution was carefully mixed with 0.45 volume of cell suspension by gently inversion. The cell suspension/working solution mixture was transferred to a 15ml conical tube then topped up to 6 ml with working solution. The working solution/cell suspension was overlaid with 3 ml of the 22% (w/v) iodixanol and the 22% iodixanol layer was overlaid with 0.5 ml of 0.04 %BSA in 1XPBS/DEPC. Viable cells were separated by density gradient created by centrifuging at 800 g for 20 min at 20°C. Viable cells were harvested from the top interface (∼500ul) and then diluted in 2 volumes (1ml) of 0.04 %BSA in 1XPBS/DEPC-treated water. The iodixanol was removed by pelleting live cell suspension at 300 g for 10 min at 4°C. Supernatant was decanted and cells were resuspended in the leftover 0.04 %BSA in 1XPBS/DEPC- treated water. Viability and concentration were measured by 0.4% trypan blue (Gibco, 15250-061) and hemocytometer. Libraries were generated with a 10X Genomics Single-cell Transcriptome Library kit.

### Bioinformatics

For purified progenitor RNAseq studies, we obtained approximately 6 million reads per biological replicate. We used FASTQC to evaluate the quality of raw, paired- end reads, and trimmed adaptors and reads of less than 36 base pairs in length from the raw reads using Trimmomatic version 0.36. We used HISAT2 version 2.1.0 to align reads to the *Drosophila* transcriptome- bdgp6, and converted the resulting BAM files to SAM flies using Samtools version 1.8. We counted converted files using Rsubread version 1.24.2 and loaded them into EdgeR. In EdgeR, we filtered genes with counts less than 1 count per million and normalized libraries for size. Normalized libraries were used to call genes that were differentially expressed among treatments. Genes with P-value < 0.05 and FDR < 0.05 were defined as differentially expressed. Principle component analysis was performed on normalized libraries using Factoextra version 1.0.5, and Gene Ontology enRIchment anaLysis and visuaLizAtion tool (GORILLA) was used to determine Gene Ontology (GO) term enrichment. Specifically, differentially expressed genes were compared in a two-list unraked comparison to all genes output from edgeR as a background set, and redundant GO terms were removed.

For single cell analysis, Cell Ranger v3.0 was used to align sequencing reads to the *Drosophila* reference transcriptome (FlyBase, r6.30) and generate feature-barcode matrices. These matrices were analyzed using the Seurat R package (version 3.2.3). Cells possessing <500 UMIs or >2500 UMIs were removed to reduce the number of low- quality cells and doublets. Seurat was then used to normalize expression values and perform integrated data cell clustering at a resolution of 0.5 with 15 principal components. Clusters were identified based on known markers and previous single-cell analysis of the *Drosophila* intestine (https://www.flyrnai.org/scRNA/). For GO term analysis of single cell data, Seurat was used to integrate *esg^ts^/+* and *esg^ts^/D30A* datasets and generate lists of differentially expressed genes for each cluster. Both up- and down-regulated gene lists (p-value cut-off <0.05) were analyzed in GOrilla to determine GO term enrichment. Differentially expressed genes were compared in a two-list unranked comparison to all genes identified in the single- cell dataset. GO terms were then analyzed in REVIGO (REduce and VIsualize Gene Ontology) to remove redundant GO terms. Top enriched GO terms are shown for each cluster, as well as those same GO terms found in other clusters. EE subset analysis was followed at Guo et al. (2019).

For Pseudotime analysis we used Monocle3 (version 0.2.0). Specifically, we converted the existing Seurat data from each genotype separately into a Monocle cell data set of midgut epithelial cells and performed trajectory analysis. We manually assigned the root node of the trajectory to the node at the tip of the Progenitor cluster for each genotype. We then subset the trajectory branch that explains pseudotime within the Progenitor population to perform all subsequent gene level analysis. Here, we manually assessed expression of genes along pseudotime with known functions in ISC identity, division, and differentiation including genes that were differentially expressed based off our Seurat analysis.

### Data availability

Gene expression data have been submitted to the NCBI GEO database (GEO: SuperSeries GSE141897 (GSE171001 and GSE141896)). Single cell gene expression data for *esg^ts^/+* flies (https://singlecell.broadinstitute.org/single_cell/study/SCP1696/single-cell-expression-data-for-d-melanogaster-wild-type-intestines) and for *esg^ts^/D30A* flies (https://singlecell.broadinstitute.org/single_cell/study/SCP1699/single-cell-expression-data-for-d-melanogaster-intestines-with-immune-deficient-progenitor-cells#study-summary) are available for visualization on the Broad Institute Single Cell Portal.

## ACKNOWLEDGEMENTS

*Drosophila* lines were kindly provided by Dr. Shelagh Campbell, Dr. Lucy O’Brien, Dr. Bruce Edgar, Dr. Heinrich Jasper and Dr. Bruno Lemaitre. We acknowledge microscopy support from Dr. Steven Ogg and Gregory Plummer at the Faculty of Medicine and Dentistry Imaging core; flow cytometry support from Dr. Aja Rieger at the Faculty of Medicine and Dentistry Flow Cytometry Core; and support with single-cell library preparation from Dr. Joaquin Lopez-Orozco. We are grateful to Dr. David Fast for assistance with analysis of the RNAseq data. The authors wish to thank Kin Chan at the Network Biology Collaborative Centre for the RNA-Seq service. Network Biology Collaborative Centre is a facility supported by Canada Foundation for Innovation, the Ontarian Government, and Genome Canada and Ontario Genomics (OGI-139). This work was supported by a grant from the Canadian Institute of Health Research (Grant # PJT 159604). Minjeong Shin was supported by Basic Science Research Program through the National Research Foundation of Korea (NRF) funded by the Ministry of Education (2020R1A6A3A0303955511). Meghan Ferguson has funding through Alberta Innovates Graduate Student Scholarships and an NSERC PGS-D.

## SUPPLEMENTARY DATA

**Figure S1:**
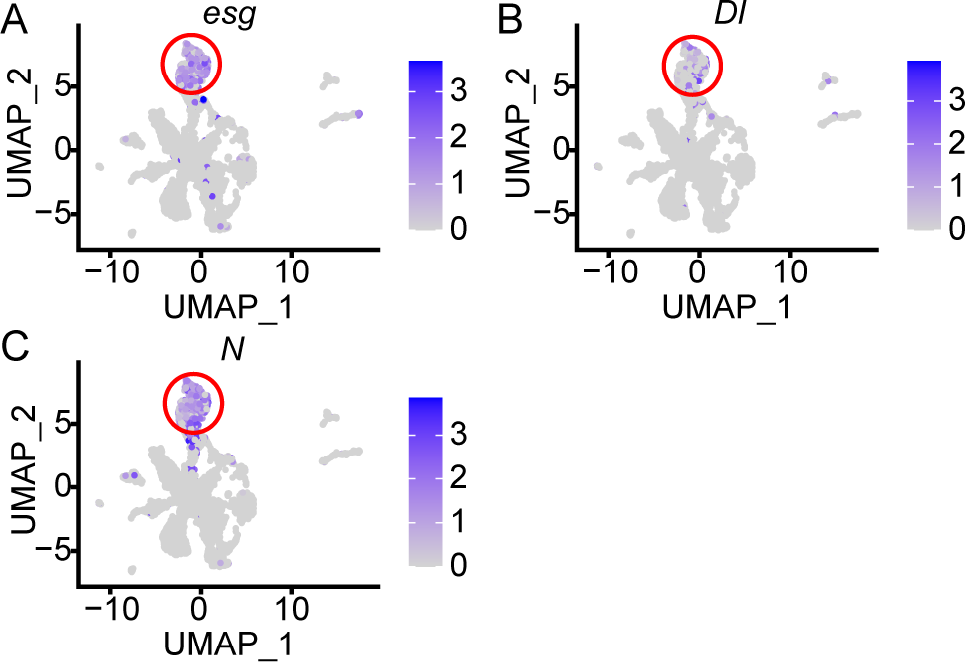
Feature plots showing expression of *Drosophila* progenitor cell markers *escargot* (*esg*), *Delta (Dl*), and *Notch* (*N*) in an unsupervised UMAP prepared with single-cell expression data from adult *esg^ts^*/*+* female *Drosophila* midguts. Progenitor cells are indicated with a red circle.

**Figure S2.**
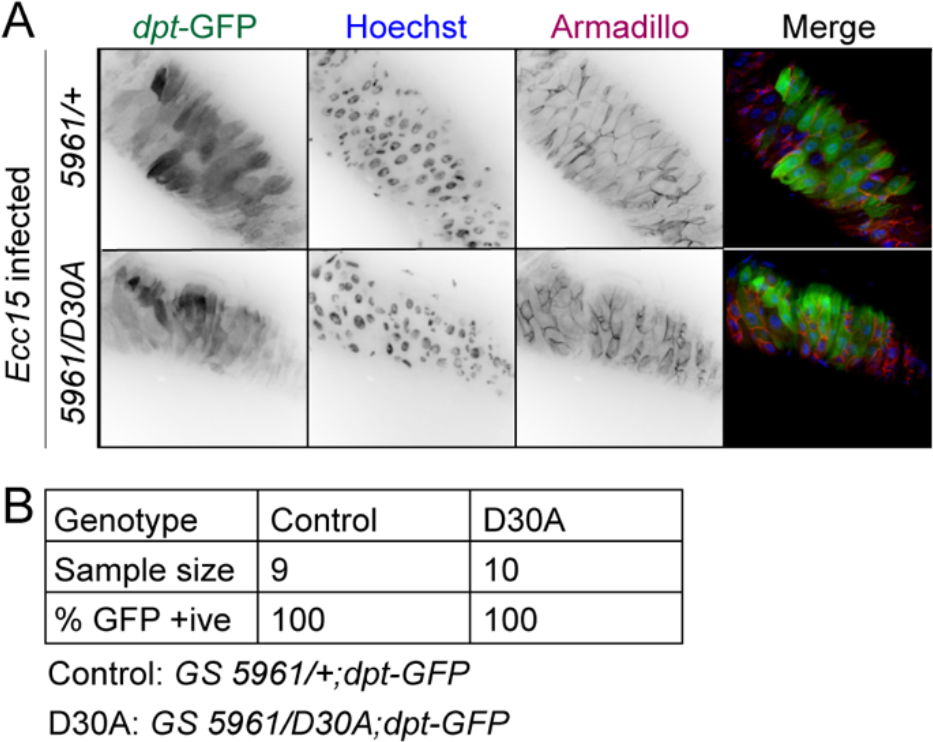
A:Visualization of the IMD pathway reporter *diptericin:GFP* (*dpt*-GFP), DNA (Hoechst), and the beta-catenin ortholog Armadillo in intestines of adult female *Drosophila* infected overnight with pathogenic *Ecc15*. The *esg^ts^* line used in the rest of the study marks progenitor cells with GFP, preventing us from unambiguously identifying cells that expressed GFP under control of the *dpt* promoter in an *esg^ts^* background. Therefore, we used the *GS5961* gene switch fly line for RU486-dependent induction of the GAL4 transcription factor in intestinal progenitor cells in this experiment. In the upper row, we visualized *dpt-*GFP expression in *GS5961/+* (*5961/+*) flies, and in the lower row, we visualized *dpt-*GFP expression in *GS5961/UAS-imdD30A* (*5961/D30A*) flies. Both lines were treated with RU486 for 48h prior to infection. **B:** Quantification of fly guts of the indicated genotypes that expressed *dpt-*GFP in enterocytes after infection with *Ecc15.* Expression of imdD30A in progenitors (*D30A*) did not prevent infection-mediated activation of IMD responses in mature enterocytes.

**Figure S3.**
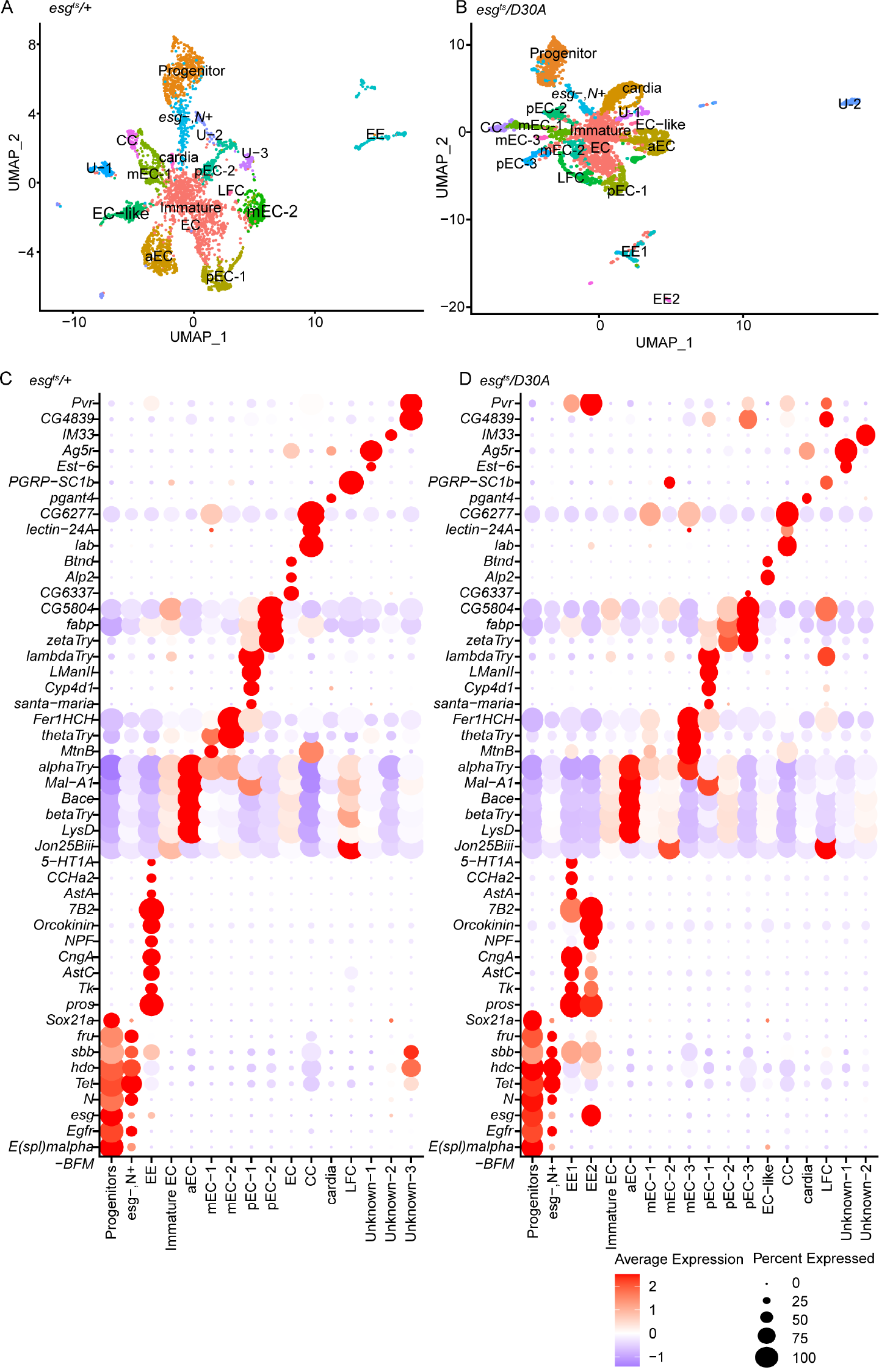
A-B: Two dimensional UMAP projection of cell types isolated from female control *esg^ts^/+* intestines, and from female *esg^ts^/D30A* intestines, color coded by cell type. U = unknown, CC = copper cells, EC = enterocyte, EE = enteroendocrine cell. **C-D:** Heatmap of IEC cluster markers colored by relative gene expression for flies of the indicated genotypes. The size of the dot indicates the proportion of cells in each cluster that expressed the indicated gene.

**Figure S4:**
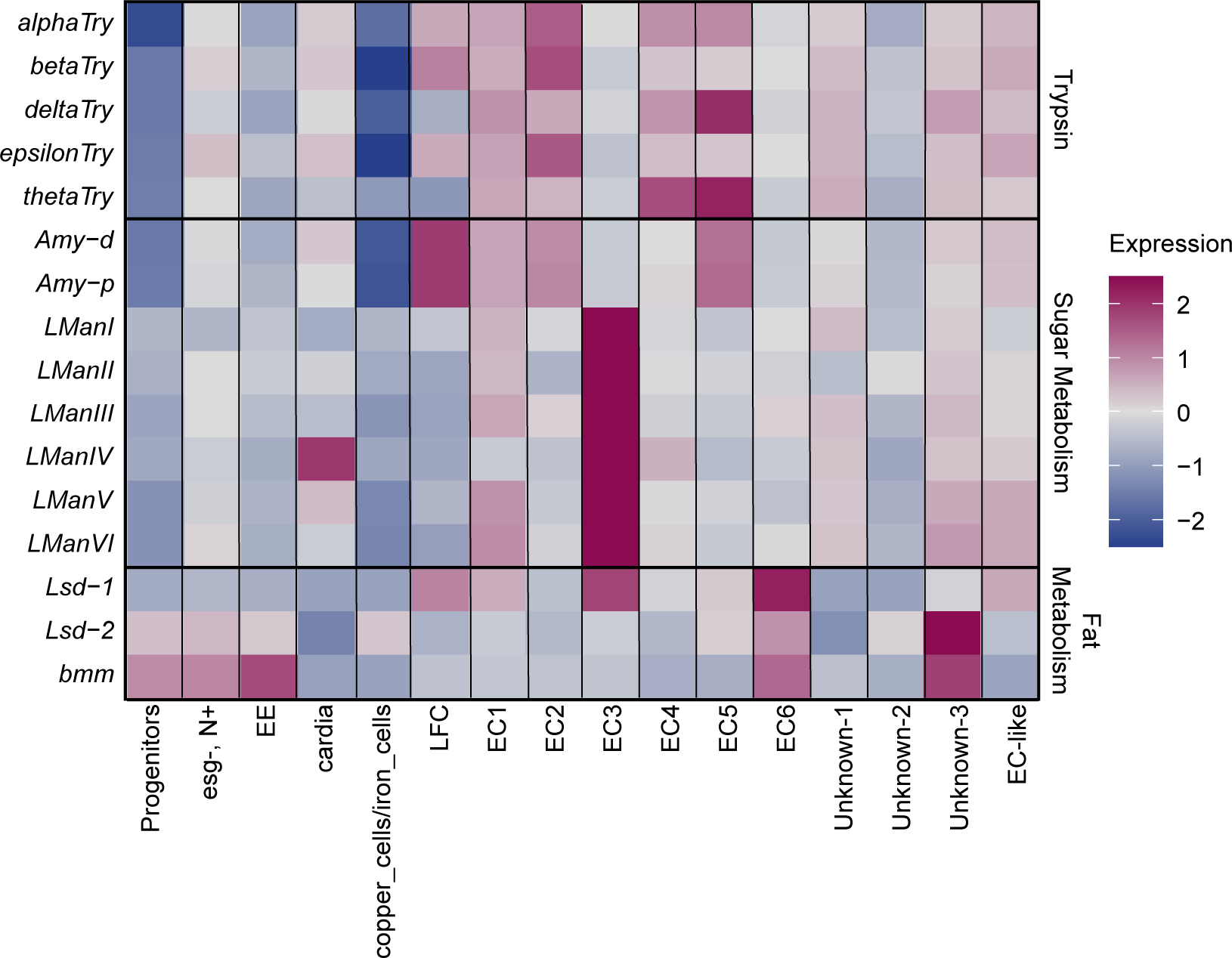
Heatmap showing relative cluster-average expression of metabolic enzymes (Try = *Trypsin*, Amy = *Amylase*, LMan = Lsysomal alpha-mannosidase, Lsd = Lipid storage droplet, bmm = *brummer*) in each intestinal epithelial cell type of control *esg^ts^/+* flies.

**Figure S5:**
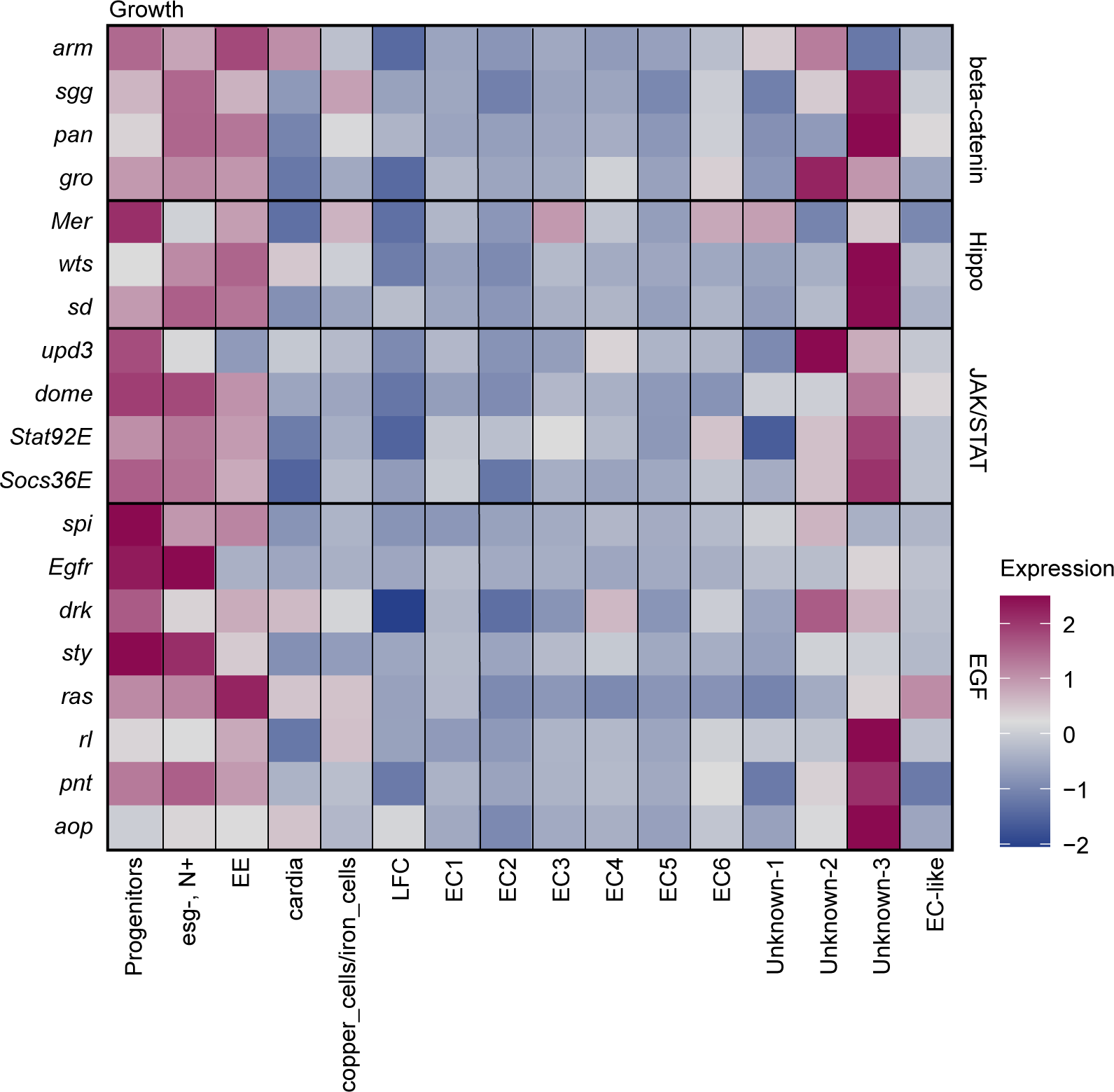
Heatmap showing relative cluster-average expression of prominent regulators of intestinal epithelial growth in each intestinal epithelial cell type of control *esg^ts^/+* flies. For this analysis, we focused on indicated components of the beta-catenin, Hippo, JAK/STAT, and Epidermal Growth Factor (EGF) pathways.

**Figure S6:**
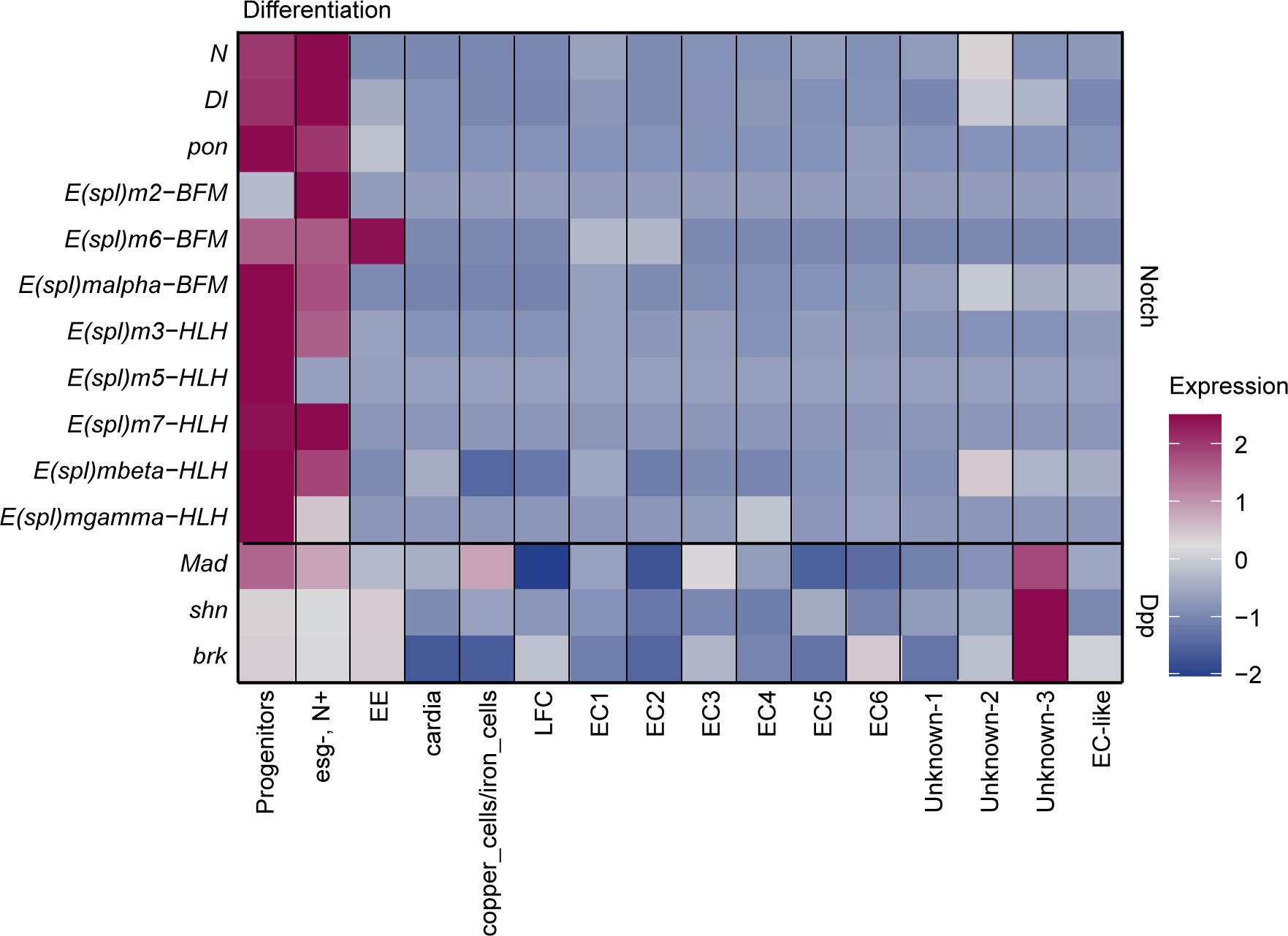
Heatmap showing relative cluster-average expression of prominent regulators of intestinal epithelial differentiation in each intestinal epithelial cell type of control *esg^ts^/+* flies. For this analysis, we focused on indicated components of the Notch and Decapentaplegic (Dpp, *Drosophila* ortholog of Bone Morphogenetic Protein) pathways.

**Figure S7:**
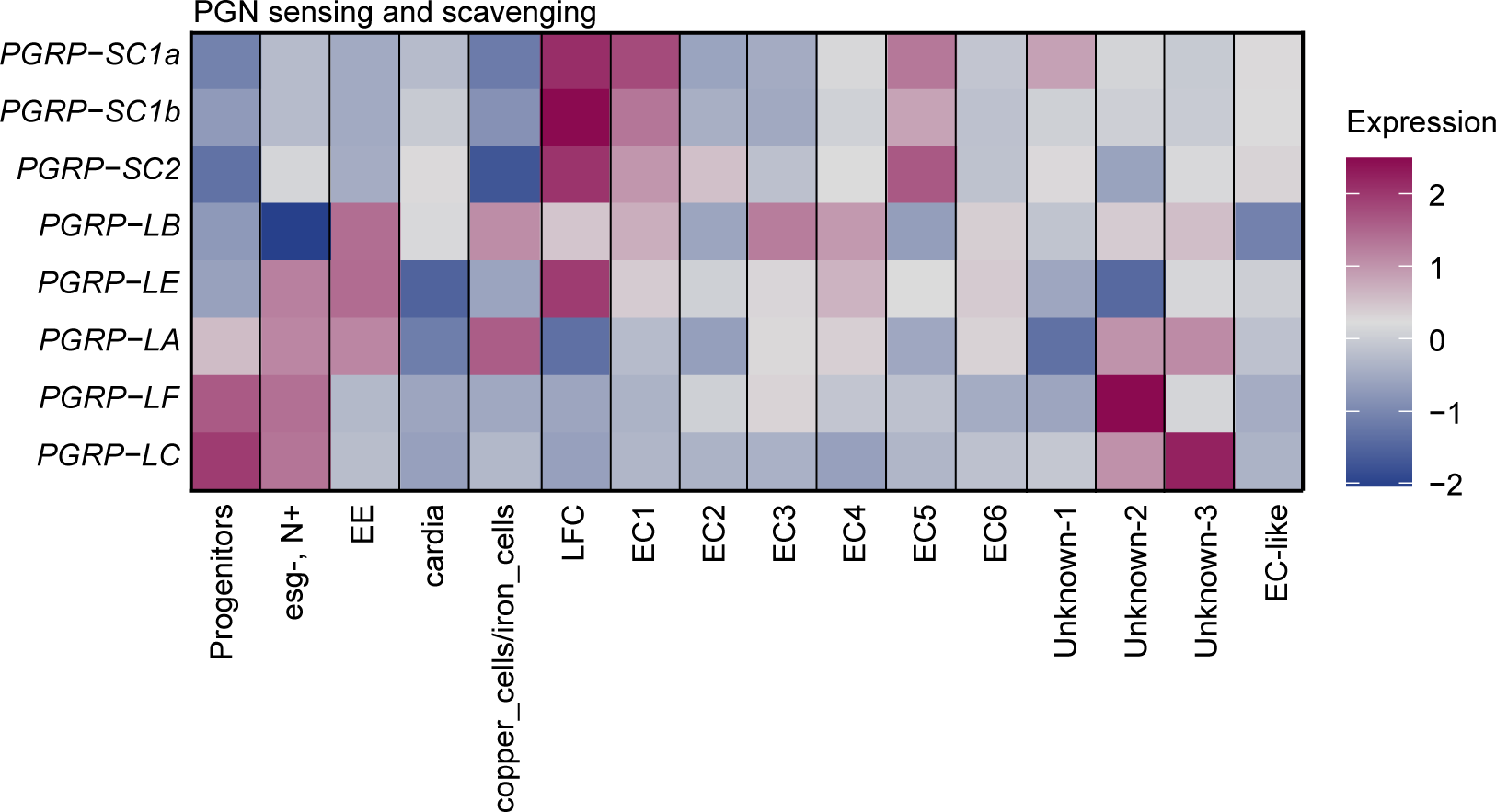
Heatmap showing relative cluster-average expression of prominent regulators of Peptidoglycan detection (*PGRP-LE, LC, LA, LF*), and peptidoglycan amidases (*PGRP-SC1a, SC1b, SC2, LB*) in each intestinal epithelial cell type of control *esg^ts^/+* flies. For this analysis, we focused on Peptidoglycan Recognition Proteins (PGRP) with detectable expression in the adult intestine.

**Figure S8:**
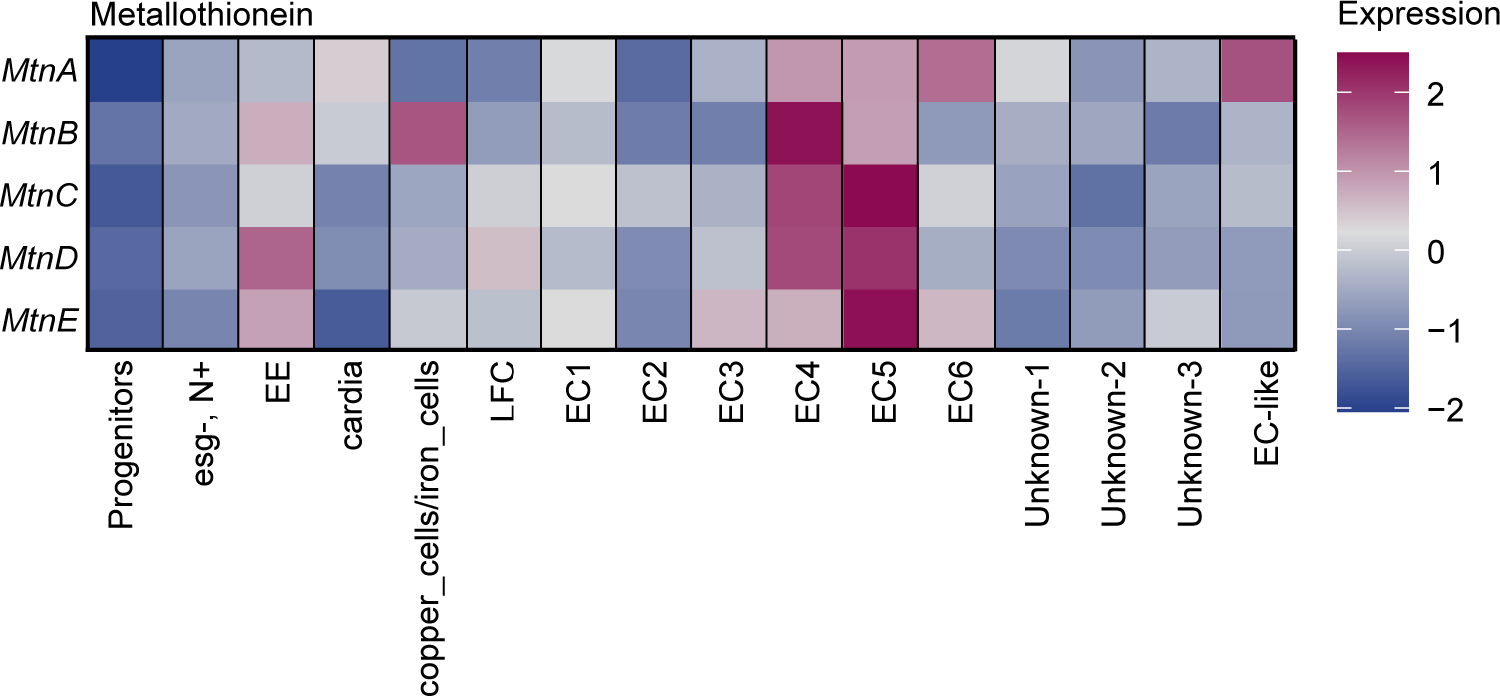
Heatmap showing relative cluster-average expression of prominent Metallothionein (*Mtn*) regulators of oxidative stress responses in each intestinal epithelial cell type of control *esg^ts^/+* flies.

**Figure S9:**
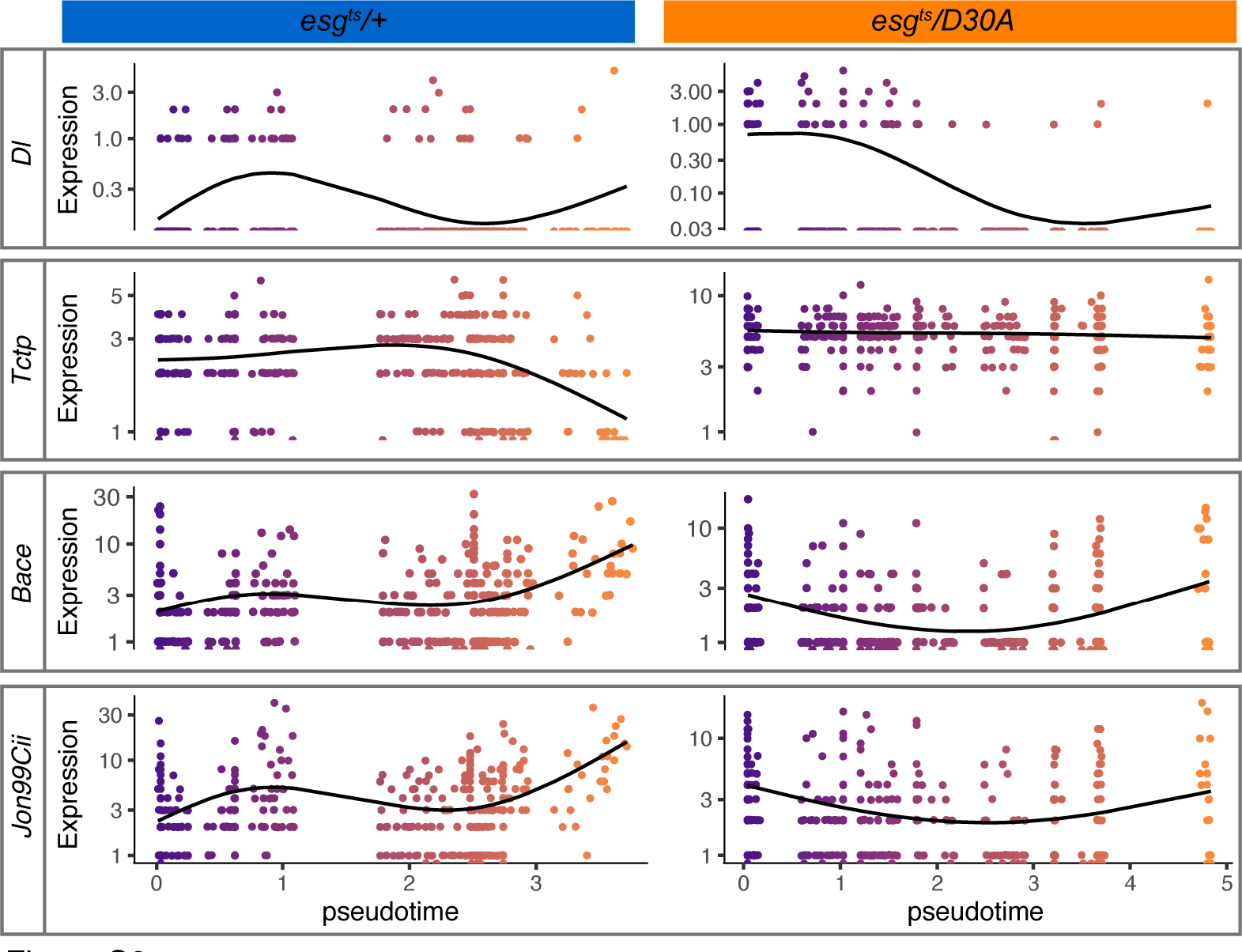
Inhibition of IMD disrupts progenitor cell expression trajectories. Expression of the stem cell marker *Delta (Dl*), *Translationally-controlled tumor protein (Tctp),* and enterocyte markers *Bace* and *Jon99Cii* along pseudotime in *esg^ts^/+* and *esg^ts^/D30A* progenitors. Dark purple marks cells at the beginning of pseudotime while orange marks cells late in pseudotime. Black lines show expression trend.

